# L-form switching confers antibiotic, phage and stress tolerance in pathogenic *Escherichia coli*

**DOI:** 10.1101/2021.06.21.449206

**Authors:** Aleksandra Petrovic Fabijan, Muhammad Kamruzzaman, David Martinez-Martin, Carola Venturini, Katarzyna Mickiewicz, Neftali Flores-Rodriguez, Jeff Errington, Jonathan R. Iredell

## Abstract

The bacterial L-form is induced by exposure to cell wall targeting antibiotics or innate immune effectors such as lysozyme and is likely to be important in many human infections. Here, we demonstrate that the osmotically fragile L-form is a distinct physiological state in Escherichia coli that is highly tolerant of oxidative stress and resistant to powerful antibiotics and common therapeutic bacteriophages. L-forms quickly revert (<20h) to their cell-walled state after antibiotic withdrawal, with apparently normal physiology and with few or no changes in DNA sequence. T4-like phages that are obligately lytic in cell-walled E. coli preferentially pseudolysogenise their L-forms providing them with transient superinfection immunity. Our data indicate that L-form switching is a common response of pathogenic E. coli strains to cell wall-targeting antibiotics and that the most commonly used lytic bacteriophages are ineffective against them in this state.

## INTRODUCTION

Bacteria proliferate in an ‘exponential phase’ under optimal conditions, e.g., when tested for antibiotic or phage susceptibility in diagnostic labs (1, 2). However, such conditions are rare in nature, where bacteria survive by adjusting their physiology and reducing their growth rates when stressed or starved (3, 4). L-forms are metabolically active bacteria that divide more slowly than the exponential phase bacteria, using a primitive mechanism that is independent of the essential FtsZ-based division machinery (5-7). Without cell walls, L-forms are osmotically sensitive and completely resistant to antibiotics targeting the cell wall (e.g., β-lactams) (5, 8). Additionally, such antibiotics often fail in the treatment of biofilm-type infections (8). These scenarios strongly invite the use of phage therapy as alternative or to complement existing cell wall-targeting antibiotics (9), which are the foundation of modern infection therapy (∼70% of all prescriptions), including sepsis management (10).

Here we demonstrate that clinical isolates of *E. coli* continue to grow without cell walls, in which state they are resistant to commonly used antibiotics (e.g., β-lactams) and highly tolerant of oxidative stress. Our newly developed methods to study phage-bacteria interactions *in vitro* demonstrate for the first time that T4-like phages that are apparently obligately lytic against normal cell-walled cells are ineffective against L-forms.

## RESULTS AND DISCUSSION

### L-form switching is a common physiological response to cell wall-targeting antibiotics in clinical *E. coli* isolates

Though first described nearly a century ago, the extent to which L-forms are an artefact of antibiotic treatment in certain bacterial species or a common adaptive response to cell wall injury remains unclear. It is also unclear whether L-forms are the result of a single process or the final endpoint of a diverse set of processes. L-forms appear to occur naturally, at least in urinary tract and biofilm-associated infections (8, 11).

Unlike non-dividing persister cells, L-forms grow and divide in the presence of powerful cell wall targeting antibiotics (5, 8), but their cell cycle and growth dynamics have not been accurately defined. To characterise L-form physiology and allow us to explore their interaction with bacteriophages, we developed an isotonic L-form growth agar (LFA) by modifying existing LFA (12) in a (i) double-layer agar plate and (ii) semisolid agar microplate format that supports the growth of cell wall-free *E. coli*. Both methods included testing of: (i) growth of typical cell-walled rods on standard hypotonic Luria-Bertani agar (positive control; LBA); (ii) growth inhibition of cell-walled forms by high-dose of cell wall-targeting antibiotics, in standard hypotonic LBA (negative control; LBA+MEM); and (iii) L-form growth induced in isotonic LFA supplemented with meropenem (LFA+MEM).

We assessed L-forms switching efficiency in 45 genetically distinct clinical *E. coli* isolates *in vitro* using meropenem, a broad-spectrum PBP2 (penicillin-binding protein 2) targeting carbapenem. This drug is currently regarded as last-line therapy for severe infections caused by multidrug resistant (MDR) *Enterobacteriaceae*, including *E. coli* (13). L-forms developed quickly in most strains tested (35/45 strains) and proliferated aerobically. The majority (85%) then quickly reverted to their normal rod-shaped cell walled forms in less than 20h after meropenem withdrawal (Fig. 1A and B). Fifteen of these strains were selected for further testing (Table 1). Revertants retained the same overall susceptibility profiles to meropenem (n=11; Table 1) as the parent precursor, with only slight variations in minimal inhibitory concentrations (MIC) in some (18%). Whole genome sequencing (WGS) on a subset of genetically distinct *E. coli* strains (Extended Fig. 1) and their revertants (n=4; Table 1) showed that at least two strains (WH62 and SYD259) were genetically identical to the strains from which they originated, with only few SNP changes in two others.

**Table 1.**
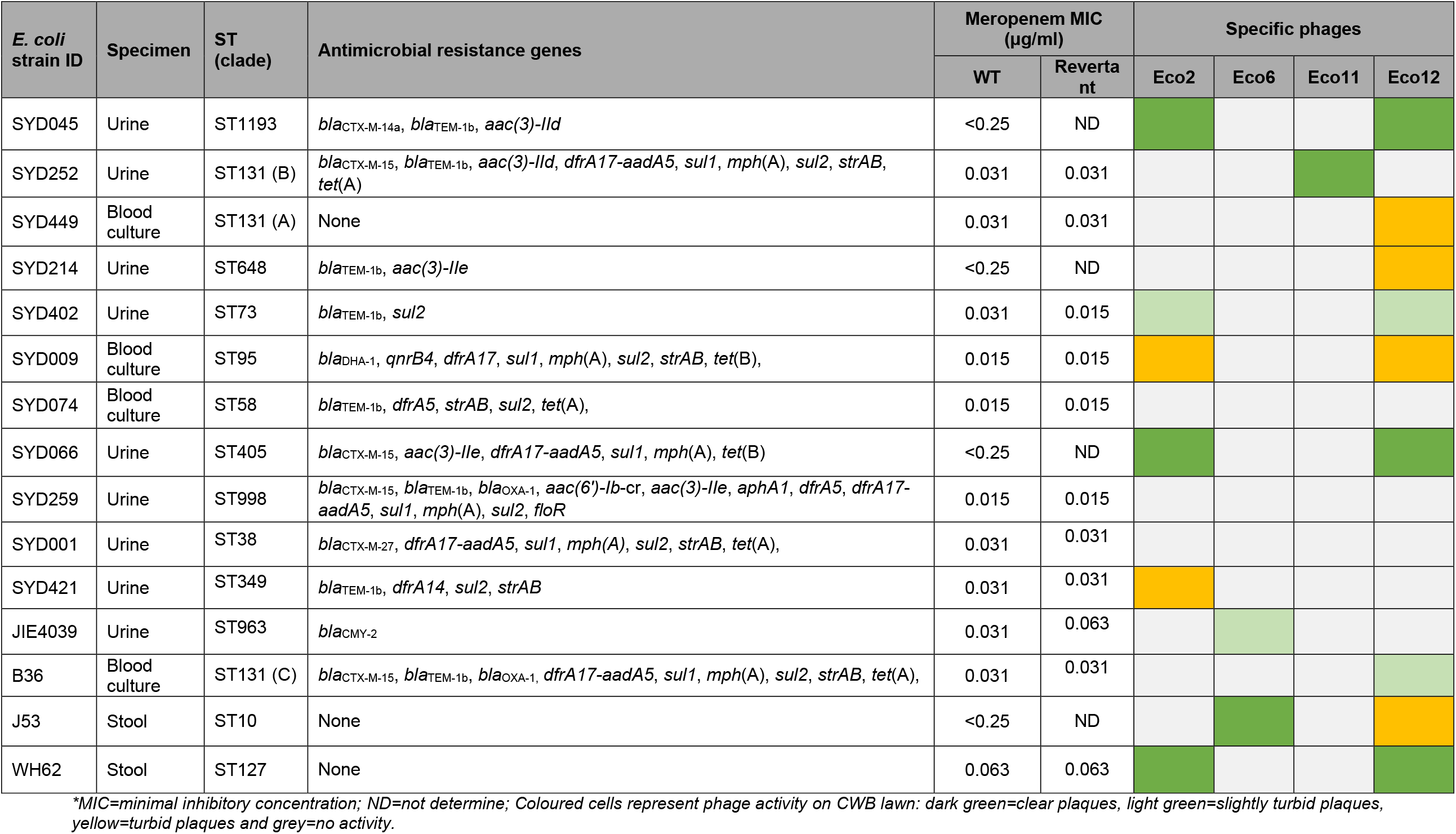
List of *E. coli* selected for further testing.

**Figure 1.**
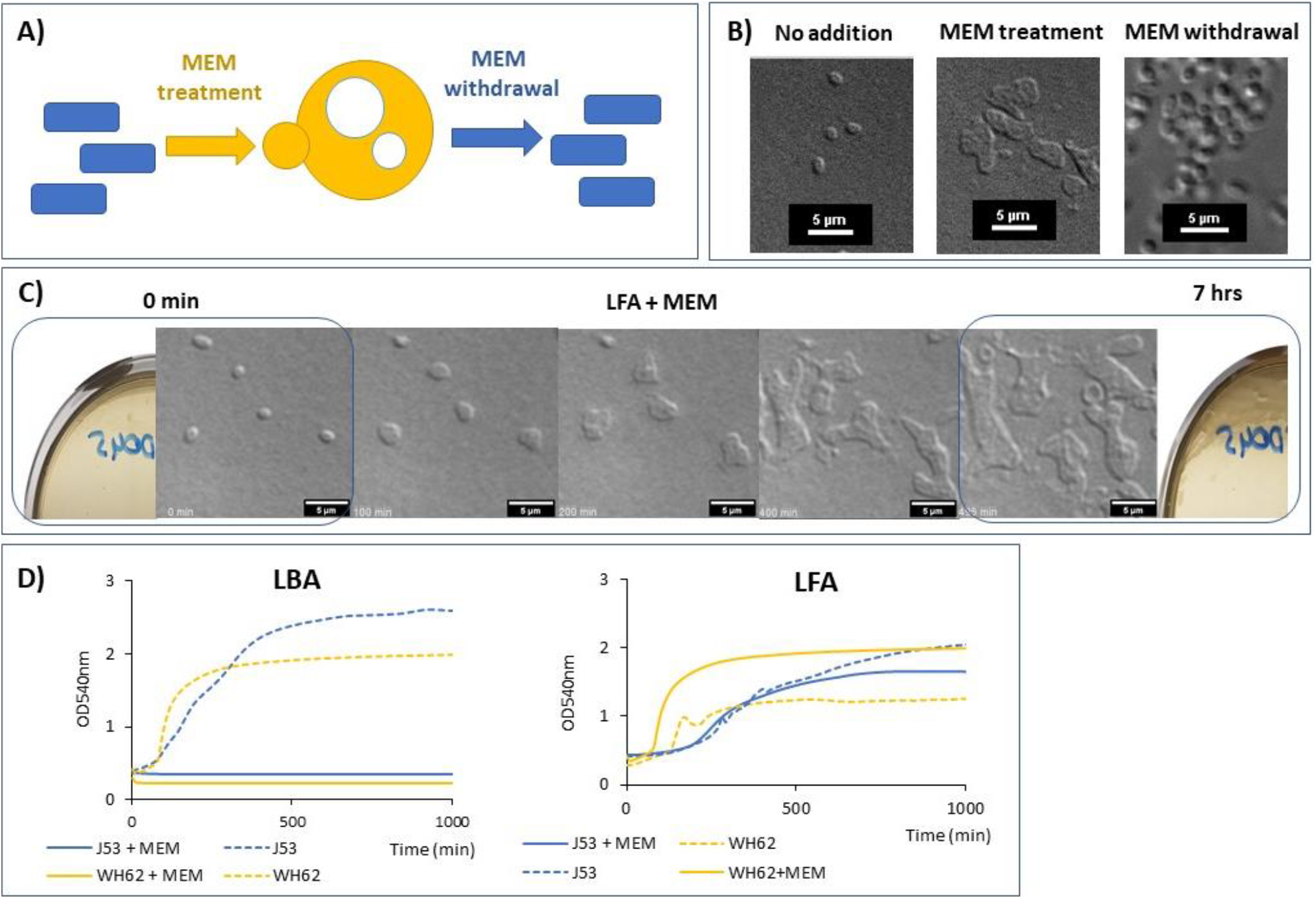
Meropenem promotes L-form growth from the cell-walled state under aerobic conditions: A) Sketch showing pathogenic *E. coli* L-forms reverting to vegetative growth as cell-walled organisms following antibiotic removal. B) *E. coli* L-form strain WH62 switch in the presence of meropenem and reversion to cell-walled state after meropenem withdrawal. C) Time-lapse DIC microscopy of WH62 L-forms; individual micrograph frames are extracted from Supplementary Movie S1. D) Growth curves of J53 (ST10) and WH62 (ST127) L-forms.

To characterise growth rate and metabolic activity of L-forms relative to walled cells, we used a semisolid microplate agar supplemented with a redox indicator (triphenyl tetrazolium chloride or TTC) and meropenem. This demonstrated that *E. coli* L-forms are less metabolically active than parent cell-walled bacteria (CWB). Plotting the growth of the two L-forms, J53 (a well-characterised K12 derivate) (14) and WH62 (an uropathogenic clinical isolate) (15) by measuring optical density at 540nm (OD540), revealed a characteristic *lag* phase followed by a slow density increase (Fig. 1D). The *lag* phase duration for WH62 and J53 was 80 and 190 minutes respectively as L-forms developed from cell-walled bacteria in osmoprotective media with meropenem, before metabolic activity increased and they began to divide (Fig. 1C). Growth rate increased after ∼200 minutes, at which stage all bacteria appeared to have converted to L-forms. After ∼500 minutes incubation, the growth rate slowed as the exponential phase ended and L-forms appeared to enter a stationary phase, reflected in reduced growth of biomass as observed under microscopy (Supplementary Movie 1) and reduced metabolic activity (Fig. 1D).

L-form growth for *E. coli* WH62 strain (ST127, a virulent subtype of uropathogenic *E. coli*) (15) was imaged using time-lapse differential interference contrast (DIC) microscopy. The addition of meropenem to exponential phase cell-walled bacteria in osmoprotective semisolid LFA led to the emergence of L-forms that differed in morphology and size (Fig. 1C). First, cells started to bulge and lose their regular shape, leading to a 4-fold (3.95 ± 1.17) increase in their surface area. L-forms underwent first division cycles after ∼5 h (Fig. 2A). Division events resembled those previously described (5, 7), with irregular shape perturbations leading to asymmetrical scission of the mother cell and procreation of a heterogeneous population of meropenem-resistant *E. coli* cells (Supplementary movie S1).

**Figure 2.**
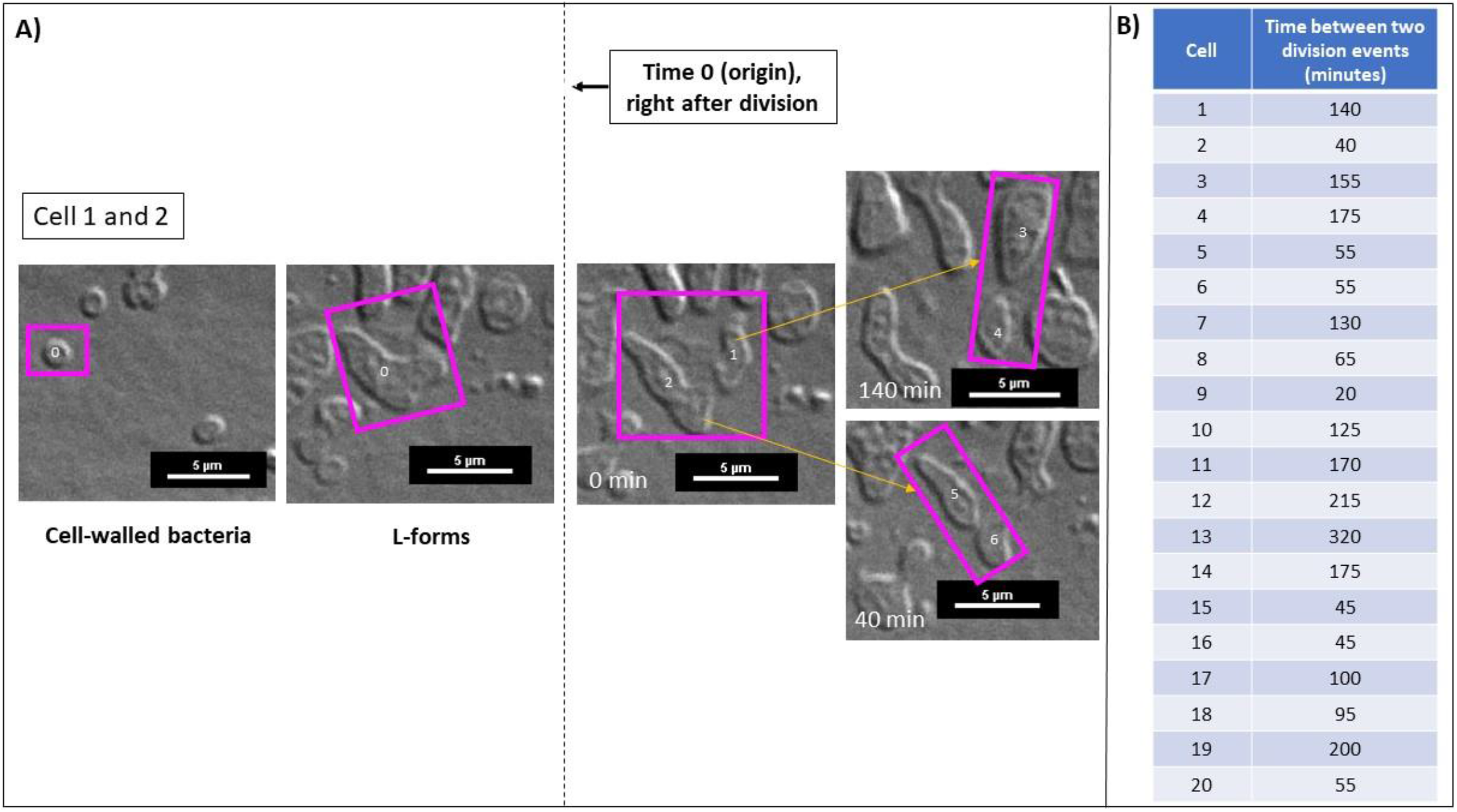
WH62 *E. coli* L-form growth and division in LFA medium with meropenem observed by time lapse DIC microscopy. A) Mode of cell division of L-forms division by budding, see also Supplementary Movie 1. Time 0 (origin) indicates the first division event in the L-form state. B) Length of L-forms cell cycle measured in 20 different L-forms cells.

We used time-lapse DIC microscopy to accurately quantify the duration of L-form proliferation cycles (Fig. 1B) at an average of 111.1 ± 76.8 minutes between consecutive division events in 20 independent cells (Fig. 2A and 2B). Intracellular vesicles were evident after prolonged incubation under aerobic conditions (>16h) (data not shown).

Altogether, these data confirm that clinical isolates of *E. coli* undergo efficient L-form switching in response to an important PBP2-targeting antibiotic in widespread use and subsequently grow and proliferate using a common ‘primordial’ mechanism (5, 16).

### L-forms of *E. coli* are relatively tolerant to oxidative stress

Under aerobic conditions, cellular levels of reactive oxygen species (ROS) are significantly increased during the walled to L-form transition, and this has been proposed to severely impair L-form growth *in vitro* (17). Kawai et al. (2015) have demonstrated that this can select for specific mutations (e.g., *ispA*^*\**^ in *Bacillus subtilis*), or a switching to anaerobic metabolism if conditions allow (e.g., *E. coli*) (17). Studies in *E. coli* L-forms, however, mostly involved testing of either experimental mutants (e.g., *ΔmurA*) (16) or stabilised L-forms (e.g., *ftsQ*^*\**^ and *mraY*^*^) (17) which could not revert to a cell-walled state without drug selection (16). Although stable L-forms appear to share similarities with natural L-forms (acquire no mutations and have the ability to revert to cell-walled state) and thus can serve as a good model to study basic L-form biology, they may be considered to be distinct and unrelated entities (18). Our data show that clinically important antibiotics (i.e., meropenem) efficiently induce L-forms in pathogenic *E. coli* isolates which efficiently proliferation aerobically.

Survival rates of meropenem-induced *E. coli* L-forms are significantly higher than that of their faster-growing cell-walled parents (Extended Fig. 2). This relatively high tolerance to oxidative stress may result from a ‘priming’ effect of ROS during L-form transition, as cell wall injury is known to trigger expression of genes involved in the oxidative stress response (17, 18). The host environment (i.e., innate immune system) not only promotes the L-form switch (18), but most likely favours L-form proliferation in the presence of otherwise inhibitory concentrations of ROS. If an L-form-like bacterial ancestor evolved before the oxygenation of the planet (19), the oxidative stress tolerance we see today may have developed subsequently as an ancient adaptation to an oxygen-enriched environment.

### Transcriptomic analysis of *E. coli* L-forms

We used RT-qPCR (20) to measure principal markers of oxidative (*elaB* and *oxyR*) and nutrient stress (*rpoS*) and DNA damage (*recA, dinB* and *lexA*) in one well-characterised laboratory strain (J53) and three widely diverse clinical strains (B36, JIE4039 and WH62) (21) (Table 1) to study the effects of L-form transition in *E. coli*.

Several genes previously reported to be upregulated in stationary phase bacteria (22) were also upregulated in L-forms (Fig. 3A). These include genes involved in general stress (*rpoS*) and oxidative stress responses (*elaB*) and, to a lesser extent, in the global response to DNA damage or the SOS response (*lexA, recA* and *dinB*). Our data also indicated possible genotype-and/or strain-specific transcriptome profiles.

**Figure 3.**
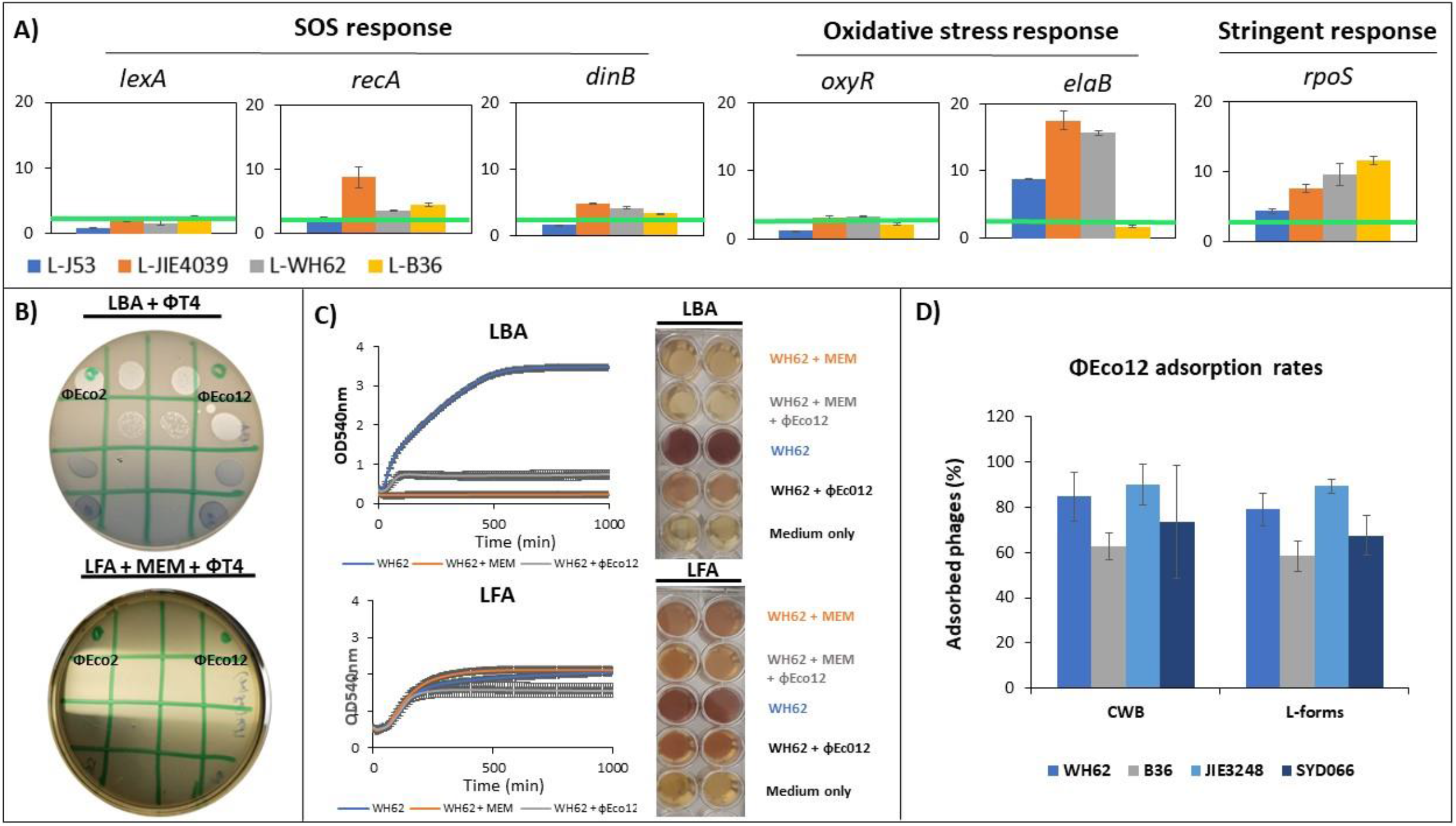
L-form transcriptomics and response to phage predation: A) Relative expression of stress response genes in *E. coli* L-forms (L-WH62, L-J53, L-B36 and L-JIE4039) in comparison with cell-walled parents (WH62, J53, B36 and JIE4039). The experiment was performed in duplicate, the mean values with standard deviations (error bars) are presented. Bright green lines represent the 2-fold difference between cell-walled bacteria and L-forms, a cut off commonly used to distinguish significant changes from insignificant ones and B) phage susceptibility of *E. coli* L-forms. WH62 meropenem-induced L-forms (bottom) displaying resistance (no lysis) to T4-like phages (Eco2 and Eco12). Control involved cell-walled counterpart on standard LBA without meropenem (top) lysed by both phages. C) Growth curves of walled bacteria (WH62 on LBA) and L-forms (L-WH62 on LFA supplemented with meropenem) in the presence of Eco12 phage (MOI 1). D) Adsorption rates (percentage of phage particles adsorbed) of Eco12 phage against walled bacteria and L-forms.

The C-tail anchored inner membrane protein ElaB was recently shown to protect bacteria against oxidative stress and heat shock and its expression to be co-ordinately regulated by both RpoS and OxyR (23). Transcription of *elaB* was strongly upregulated in three of four tested L-forms and appeared to be associated with upregulation of *rpoS* but not *oxyR* (in which no significant changes were seen) (23). However, B36 *E. coli* displayed relatively high tolerance of sublethal H2O2 in the cell-walled state and *elaB* expression changed little after L-form transition, suggesting possible alternative protective pathways in this notoriously pathogenic subtype (ST131, clade C). Consistent with previous findings (17), our results suggest that oxidative stress supports L-form proliferation in otherwise vulnerable *E. coli*.

Oxidative stress is also a mediator of DNA damage in bacteria and an important exogenous trigger of the SOS response. We find that transcription of the low-fidelity DNA polymerase Pol IV (*dinB*) (24) (early SOS response) and the repair protein gene *recA* (late SOS response) (25) are slightly upregulated in all *E. coli* L-forms we tested. Interestingly, the level of *recA* transcripts detected in L-forms of JIE4039 (ST963) were about 7-fold higher than the level found in other three L-forms, suggesting that this particular genotype might be more sensitive to oxidative stress than others. Finally, induction of the master SOS response regulator *lexA* did not differ significantly between the walled and L-forms of the bacteria tested, suggesting that *E. coli* L-forms did not suffer extensive DNA damage when exposed to sublethal ROS.

We also found highly consistent upregulation (twofold or greater) of *rpoS* after the transition to L-form. Translation of *rpoS* and induction of the stringent response mediated by expression of specific snRNAs (e.g., *oxyS* and *dsrA*) under different stress conditions (26) is well described in stationary phase bacteria and in biofilms (27). Our data suggest that RpoS-regulated genes and gene pathways, including those involved in DNA repair and oxidative stress responses, may be responsible at least in part for the increase in membrane resilience and high oxidative tolerance observed in the *E. coli* L-forms studied here.

### L-forms of *E. coli* resist common phages

Very little is known about the interaction between predatory bacteriophages and L-forms, but this is a bacterial stress that is as ancient as ROS. Phage therapy is often advocated for recurrent and biofilm-associated infections (28) but there is a little experimental evidence to support this. In fact, certain therapeutic phages were shown to fail *in vivo* despite their obvious effective killing observed *in vitro* (29) and this may have resulted in the failure of the largest prospective, randomized, placebo-controlled, parallel-group clinical trial (30). We therefore tested various obligately lytic phages that are obvious candidates for use in phage therapy against L-forms. Surprisingly, we found that vB_EcoM_2 (Eco2) and vb_EcoM_12 Eco12 *E. coli* specific phages, (Fig. 3B and Extended Fig. 3) characterised here as T4-like (fam. *Myoviridae*) fail to propagate in L-forms of bacterial strains against which they are powerfully lytic of cell-walled forms (CWB) *in vitro*. Phage adsorption assays indicate that T4-like phages adsorb to L-forms with similar efficiency to CWB (Fig. 3D) but a plaque assay revealed no lysis whatsoever of the L-form lawn (Fig. 3B). L-form growth curves are also unaffected by the presence of phages (Fig. 3C) which lyse their CWB parents. Pulsed Field Gel Electrophoresis (PFGE) revealed an extra DNA fragment corresponding in size to the T4-phage genomes, which are resistant to digestion by *Xba1* or *Apa1* (which are sensitive to restriction site methylation by Dam and Dcm, respectively). Taken together, these data indicate preferential pseudolysogeny in *E. coli* L-forms with persistence of these phages within L-forms as methylated episomes (Extended Fig. 4). Genome annotation of Eco12 phage identifies genes previously been implicated in pseudolysogeny (31-33), including a superinfection exclusion gene (gp_140), putative *rI* lysis inhibition gene (gp_204) and methyltransferase genes (gp_125 and gp_138) (31). The immunity provided by the episomal phage during pseudolysogeny has previously been reported to allow survival and subsequent emergence of phage resistant mutants (31), which can cause therapeutic failure during phage therapy.

The common indications for phage therapy include urinary tract and other infections in which conditions are likely to favour L-forms (8) and in which powerful cell wall-active drugs are usually co-administered. Our findings should be considered a strong note of caution when considering co-administration of antibiotics such as meropenem with common T4-like phages, as this appears likely to result in simultaneous and apparently complete failure of antibiotic and phage therapy when it is least desirable. It is important to carefully consider both bacterial and phage physiology before embarking on phage therapy.

## METHODS

### Bacterial strains and DNA sequencing

The potential for L-from growth was tested in a wide range of *E. coli* strains (n=45), including a multidrug resistant dominant clone ST131. The testing also involved strains belonging to other clinically important STs such as: 648, 63, 224, 69, 193, 12, 127, 349, 62, 73, 95, 05, 58, 80, 998, 38, 453 and 10. Table 1 lists the genetically distinct strains (n=15) selected for further testing which included testing of meropenem susceptibility in revertants and phage-susceptibility. DNA from overnight cultures of revertants was extracted using a DNeasy blood and tissue kit (Qiagen). Paired-end multiplex libraries were prepared for *E. coli* isolates using the Illumina Nextera XT kit and sequenced on the Illumina NextSeq 500 NCS v2.0 platform. Reads were quality-checked, trimmed and assembled using publicly available tools, including SPAdes 3.9.0 (*de novo*) and Progressive Mauve (34). Single-nucleotide polymorphisms were identified using snippy v.3.1 (https://github.com/tseemann/snippy). Sequence typing was performed using mlst (*in silico* multilocus sequence typing) (https://github.com/tseemann/mlst). Read mapping for revertant genomes was performed against the assembled, reordered sequence of the original cell-walled strains.

### Growth conditions

*E. coli* isolates were grown on Brilliance GBS Agar/Oxoid (Thermo Fisher Scientific) and in Luria-Bertani broth. Bacterial L-forms were grown in osmoprotective L-form medium (LFA) as described previously (12). When necessary, antibiotics and supplements were added at the following concentrations: meropenem (100μg/ml) and 2,3,5-triphenyl-2H-tetrazolium chloride or TTC (5%). Reversion to the cell wall state was demonstrated by plating out L-forms on both LFA and LBA (<10% survived) without antibiotics.

### Antibiotic susceptibility assay

MICs of meropenem in revertans was assessed using microbroth-dilution protocol as previously described and interpreted according to Clinical and Laboratory Standard Institute (1).

### Microscopic imaging and growth kinetics

Sample preparation for time-lapse DIC microscopy has been done as previously described with a slight modification (35). Instead of standard imaging medium, LFA supplemented with meropenem was used. DIC microscopy images were acquired at 37°C on a Nikon Eclipse Ti-E motorised inverted microscope with Perfect Focus using a Nikon 100x 1.45 N.A PlanApo Lambda objective (Nikon Instruments). Images were captured using a Nikon DS-Qi2 monochrome camera at 5 min intervals for up to 16 h using the NIS-Elements software (Laboratory Imaging s.r.o.). DIC illumination was achieved using Nomarski prisms. Pictures and videos were prepared for publication using ImageJ (http://rsb.info.nih.gov/ij).

### Hydrogen peroxide assay and L-form transcriptome

Tolerance to H2O2 were determined by exposing exponential phase cell-walled and L-form bacteria at a density of ∼2×108 CFU/ml to sublethal concentrations (10 mM) of hydrogen-peroxide. Bacterial cultures were sampled at different time intervals (0, 20, and 60 min) and dilutions were spread onto LB agar (cell-walled bacteria) and LFA (L-forms) plates and grown at 37 °C for a further 18 hrs. To determine the stress responses, L-forms were induced as described above and RNA extraction, cDNA preparation and qRT-PCR were performed as described elsewhere (20). Supplementary table 1 contains the primers used in RT-PCR assay to define L-form stress responses.

### Isolation and genome sequencing of Escherichia coli specific phages

*Bacteriophages Eco2 and Eco12 targeting pathogenic E. coli were isolated from sewage and pond water samples respectively collected in the Greater Sydney District (Sydney, NSW, Australia) during 2019. Specimens were clarified by filtration through a 0*.*45 μm and 0*.*22 μm filters. Isolation of bacteriophages was performed using an enrichment procedure (36) where single plaques were picked and purified as previously described (37). High-titer stocks were prepared by propagating bacteriophages over several double-layer plates washed in SM buffer (50 mM Tris-HCl, 8 mM MgSO4, 100 mM NaCl, pH 7*.*4), filtered through a 0*.*22 μm filter and precipitated with NaCl and PEG8000 (37). The concentration as plaque forming units per mL (PFU/mL) was determined by spotting 10 μL of 10-fold serial dilutions onto a double-layer of the target bacteria (37). High-titer (*≥*1010 PFU/mL) bacteriophage stocks were stored at 4°C. Bacteriophage DNA isolation, sequencing and genome assembly were performed as described previously (38)*.

### Phage susceptibility and phage adsorption assay

Phage-susceptibility testing was performed using a traditional plaque or a double-layer agar method as previously described (39). When testing phage-susceptibility in L-forms, instead of standard LBA medium, LFA supplemented with meropenem was used. Inhibition of cell-walled and L-form bacterial growth was determined as described previously (40) with modification that included LFA supplemented with meropenem to support L-form growth. The adsorption of phage to cell-walled and L-form bacteria was determined by a method described elsewhere (40). Briefly, the phages and bacterial suspensions were mixed in SM buffer (cell-walled bacteria) and L-form broth (L-forms) at a multiplicity of infection (MOI) 0.01 and incubated at various temperatures for 30 min. The mixtures were centrifuged (10,000 x g, 15 min), and unabsorbed phage counts in supernatant were determined. Phage adsorption rates were expressed as percentages of adsorbed phages in relation to the initial phage counts. The data were plotted and fitted with exponential curves using Excel Version 2105.

### Pulsed Field Electrophoresis Gel (PFGE)

The presence of episomal phage in WH62 L-form isolate was determined using the PFGE method described elsewhere (41). Briefly, freshly induced WH62 L-form was co-incubated with Eco12 phage during 3hrs with continuous shaking (225rpm) at 37 °C. After incubation, infected cells were washed three times with L-form broth (2500 x g, 10 min) in order to remove extracellular phages. In order to determine methylation of episomal phages, the DNA was digested with S1-nuclease (42). Electrophoresis was carried out for 20 h at 14 °C, pulse times 6-36 s at 6 V/cm on a Bio-Rad CHEF MAPPER apparatus (Bio-Rad Laboratories). Restriction profiles were analyzed using the BioNumerics version 7.10 finger-printing software.

## Data availability

All data generated or analysed during this study are included in this article and its Supplementary Material. Whole genome sequencing data are available on NCBI under the BioProject accession number PRJNAXXXXX.

**Supplementary Movie 1**. *E. coli* (WH62) transition from rod to L-form on osmoprotective medium supplemented with meropenem (100μg/ml). DIC images were acquired automatically every 5 min for about 5 hrs. Scale bar, 5 μm.

## Acknowledgments

We are grateful to Alicia Fajardo Lubian and Sally Partridge for their advice on experimental design. We thank Dr. Nouri Ben Zakour from Westmead Institute for Medical Research for her help with phage genome sequencing data and staff at the Pathogen Genomics Unit, Westmead Hospital for their technical advice and sequencing support. The authors acknowledge the technical and scientific assistance of Sydney Microscopy and Microanalysis, the University of Sydney node of Microscopy Australia. This work was funded by National Health Medical Research Council (Australian Government) Investigator Grant (Iredell_APP1197534).

## Author contributions

A.P.F. and J.R.I. conceived the study and designed the main experimental plan. A.P.F., J.R.I., J.E. and K.M. analysed the data. A.P.F. and J.R.I. wrote the paper. A.P.F. developed double-layer plaque and microtiter assay that support the L-forms growth *in vitro*. A.P.F., D.M.M. and N.F.R. designed the L-form microscopy experiments. M.K. and A.P.F. designed transcriptome experiments. C.V. analysed WGS data of *E. coli* isolates and revertants. A.P.F. performed all experiments. All authors were involved in reviewing and editing the final manuscript.

## Competing interest

The authors declare no competing interest.

## Notes

### Competing Interest Statement

The authors have declared no competing interest.

## References

1. Performance Standards for Antimicrobial Susceptibility Testing, 31st Edition 2021 [Available from: https://clsi.org/standards/products/microbiology/documents/m100/.

2. Petrovic Fabijan A, Lin RCY, Ho J, Maddocks S, Ben Zakour NL, Iredell JR, et al. Publisher Correction: Safety of bacteriophage therapy in severe Staphylococcus aureus infection. Nat Microbiol. 2020;5(4):652.

3. Poole K. Bacterial stress responses as determinants of antimicrobial resistance. J Antimicrob Chemoth. 2012;67(9):2069–89.

4. Kolter R, Siegele DA, Tormo A. The stationary phase of the bacterial life cycle. Annu Rev Microbiol. 1993;47:855–74.

5. Mercier R, Kawai Y, Errington J. General principles for the formation and proliferation of a wall-free (L-form) state in bacteria. Elife. 2014;3.

6. Dai K, Lutkenhaus J. Ftsz Is an Essential Cell-Division Gene in Escherichia-Coli. J Bacteriol. 1991;173(11):3500–6.

7. Mercier R, Kawai Y, Errington J. Excess membrane synthesis drives a primitive mode of cell proliferation. Cell. 2013;152(5):997–1007.

8. Mickiewicz KM, Kawai Y, Drage L, Gomes MC, Davison F, Pickard R, et al. Possible role of L-form switching in recurrent urinary tract infection. Nat Commun. 2019;10(1):4379.

9. Petrovic Fabijan A, Lin RCY, Ho J, Maddocks S, Ben Zakour NL, Iredell JR, et al. Safety of bacteriophage therapy in severe Staphylococcus aureus infection. Nat Microbiol. 2020;5(3):465–72.

10. Abdul-Aziz MH, Sulaiman H, Mat-Nor MB, Rai V, Wong KK, Hasan MS, et al. Beta-Lactam Infusion in Severe Sepsis (BLISS): a prospective, two-centre, open-labelled randomised controlled trial of continuous versus intermittent beta-lactam infusion in critically ill patients with severe sepsis. Intens Care Med. 2016;42(10):1535–45.

11. Blango MG, Mulvey MA. Persistence of uropathogenic Escherichia coli in the face of multiple antibiotics. Antimicrob Agents Chemother. 2010;54(5):1855–63.

12. Davison F, Chapman J, Mickiewicz K. Isolation of L-form Bacteria from Urine using Filtration Method. Jove-J Vis Exp. 2020(160).

13. Papp-Wallace KM, Endimiani A, Taracila MA, Bonomo RA. Carbapenems: past, present, and future. Antimicrob Agents Chemother. 2011;55(11):4943–60.

14. Yi H, Cho YJ, Yong D, Chun J. Genome sequence of Escherichia coli J53, a reference strain for genetic studies. J Bacteriol. 2012;194(14):3742–3.

15. Li D, Reid CJ, Kudinha T, Jarocki VM, Djordjevic SP. Genomic analysis of trimethoprim-resistant extraintestinal pathogenic Escherichia coli and recurrent urinary tract infections. Microb Genom. 2020;6(12).

16. Studer P, Staubli T, Wieser N, Wolf P, Schuppler M, Loessner MJ. Proliferation of Listeria monocytogenes L-form cells by formation of internal and external vesicles. Nature Communications. 2016;7.

17. Kawai Y, Mercier R, Wu LJ, Dominguez-Cuevas P, Oshima T, Errington J. Cell Growth of Wall-Free L-Form Bacteria Is Limited by Oxidative Damage. Curr Biol. 2015;25(12):1613–8.

18. Glover WA, Yang YQ, Zhang Y. Insights into the Molecular Basis of L-Form Formation and Survival in Escherichia coli. Plos One. 2009;4(10).

19. Errington J, Mickiewicz K, Kawai Y, Wu LJ. L-form bacteria, chronic diseases and the origins of life. Philos Trans R Soc Lond B Biol Sci. 2016;371(1707).

20. Kamruzzaman M, Iredell J. A ParDE-family toxin antitoxin system in major resistance plasmids of Enterobacteriaceae confers antibiotic and heat tolerance. Sci Rep. 2019;9(1):9872.

21. Riley LW. Pandemic lineages of extraintestinal pathogenic Escherichia coli. Clin Microbiol Infec. 2014;20(5):380–90.

22. Navarro Llorens JM, Tormo A, Martinez-Garcia E. Stationary phase in gram-negative bacteria. FEMS Microbiol Rev. 2010;34(4):476–95.

23. Guo YX, Li YM, Zhan WE, Wood TK, Wang XX. Resistance to oxidative stress by inner membrane protein ElaB is regulated by OxyR and RpoS. Microb Biotechnol. 2019;12(2):392–404.

24. Wagner J, Gruz P, Kim SR, Yamada M, Matsui K, Fuchs RP, et al. The dinB gene encodes a novel E. coli DNA polymerase, DNA pol IV, involved in mutagenesis. Mol Cell. 1999;4(2):281–6.

25. Little JW, Edmiston SH, Pacelli LZ, Mount DW. Cleavage of the Escherichia coli lexA protein by the recA protease. Proc Natl Acad Sci U S A. 1980;77(6):3225–9.

26. Battesti A, Majdalani N, Gottesman S. The RpoS-mediated general stress response in Escherichia coli. Annu Rev Microbiol. 2011;65:189–213.

27. Jaishankar J, Srivastava P. Molecular Basis of Stationary Phase Survival and Applications. Front Microbiol. 2017;8:2000.

28. Doub JB. Bacteriophage Therapy for Clinical Biofilm Infections: Parameters That Influence Treatment Protocols and Current Treatment Approaches. Antibiotics (Basel). 2020;9(11).

29. Aslam S, Lampley E, Wooten D, Karris M, Benson C, Strathdee S, et al. Lessons Learned From the First 10 Consecutive Cases of Intravenous Bacteriophage Therapy to Treat Multidrug-Resistant Bacterial Infections at a Single Center in the United States. Open Forum Infect Dis. 2020;7(9):ofaa389.

30. Sarker SA, Sultana S, Reuteler G, Moine D, Descombes P, Charton F, et al. Oral Phage Therapy of Acute Bacterial Diarrhea With Two Coliphage Preparations: A Randomized Trial in Children From Bangladesh. EBioMedicine. 2016;4:124–37.

31. Los M, Wegrzyn G, Neubauer P. A role for bacteriophage T4 rI gene function in the control of phage development during pseudolysogeny and in slowly growing host cells. Res Microbiol. 2003;154(8):547–52.

32. Brathwaite KJ, Siringan P, Connerton PL, Connerton IF. Host adaption to the bacteriophage carrier state of Campylobacter jejuni. Res Microbiol. 2015;166(6):504–15.

33. Siringan P, Connerton PL, Cummings NJ, Connerton IF. Alternative bacteriophage life cycles: the carrier state of Campylobacter jejuni. Open Biol. 2014;4(3).

34. Darling ACE, Mau B, Blattner FR, Perna NT. Mauve: Multiple alignment of conserved genomic sequence with rearrangements. Genome Res. 2004;14(7):1394–403.

35. de Jong IG, Beilharz K, Kuipers OP, Veening JW. Live Cell Imaging of Bacillus subtilis and Streptococcus pneumoniae using Automated Time-lapse Microscopy. Jove-J Vis Exp. 2011(53).

36. Knezevic P, Kostanjsek R, Obreht D, Petrovic O. Isolation of Pseudomonas aeruginosa specific phages with broad activity spectra. Curr Microbiol. 2009;59(2):173–80.

37. Petrovic A, Kostanjsek R, Rakhely G, Knezevic P. The First Siphoviridae Family Bacteriophages Infecting Bordetella bronchiseptica Isolated from Environment. Microb Ecol. 2017;73(2):368–77.

38. Venturini C, Ben Zakour NL, Bowring B, Morales S, Cole R, Kovach Z, et al. Fine capsule variation affects bacteriophage susceptibility in Klebsiella pneumoniae ST258. FASEB J. 2020;34(8):10801–17.

39. Anderson B, Rashid MH, Carter C, Pasternack G, Rajanna C, Revazishvili T, et al. Enumeration of bacteriophage particles: Comparative analysis of the traditional plaque assay and real-time QPCR-and nanosight-based assays. Bacteriophage. 2011;1(2):86–93.

40. Knezevic P, Obreht D, Curcin S, Petrusic M, Aleksic V, Kostanjsek R, et al. Phages of Pseudomonas aeruginosa: response to environmental factors and in vitro ability to inhibit bacterial growth and biofilm formation. J Appl Microbiol. 2011;111(1):245–54.

41. Han H, Zhou H, Li H, Gao Y, Lu Z, Hu K, et al. Optimization of pulse-field gel electrophoresis for subtyping of Klebsiella pneumoniae. Int J Environ Res Public Health. 2013;10(7):2720–31.

42. Barton BM, Harding GP, Zuccarelli AJ. A General-Method for Detecting and Sizing Large Plasmids. Anal Biochem. 1995;226(2):235–40.

