## Supplementary Data for "L-form switching confers antibiotic, phage and stress tolerance in pathogenic *Escherichia coli*"

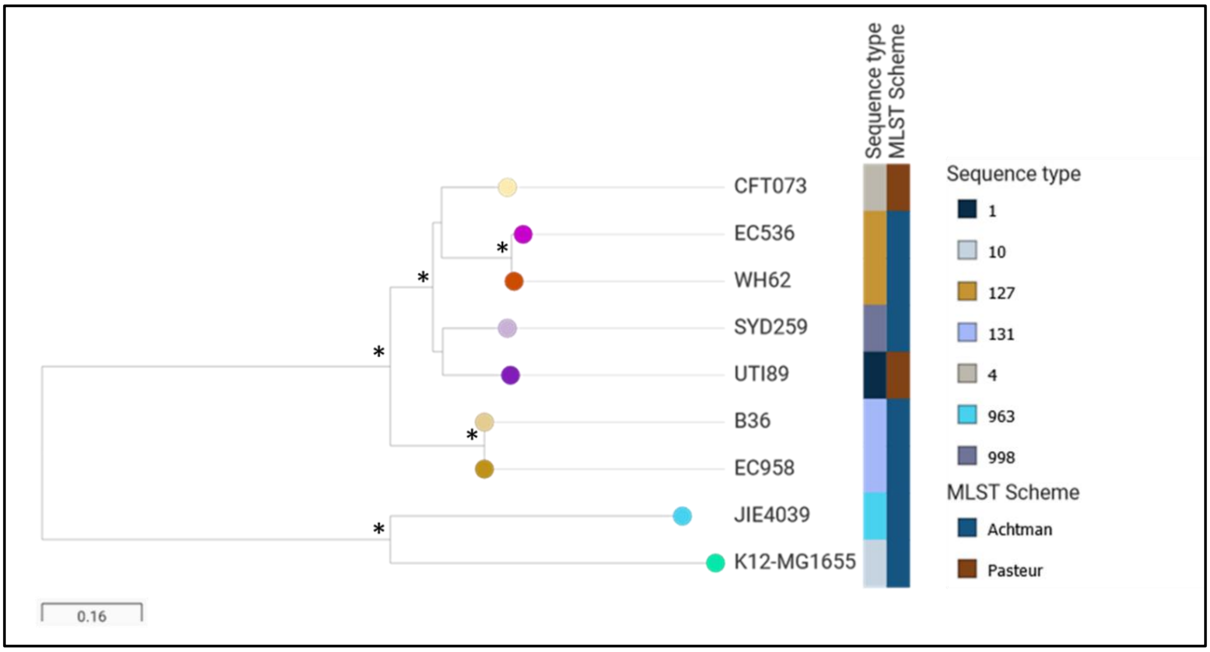


**Extended Figure 1**. Phylogenetic relationship of sequenced *E. coli* revertant strains. Maximum-likelihood phylogram showing the relatedness of *E. coli* B36 (GenBank NZ_LR130545), JIE4039 (GenBank MS14384), SYD259 (sequenced for this study) and WH62 (sequenced for this study) from our clinical collections with representative genomes of extraintestinal pathogenic *E. coli* (38): CFT073 (Genbank NC_004431), UTI89 (GenBank NC_007946), EC958 (GenBank HG941718) and *E. coli* isolate 536 (GenBank CP000247). Single nucleotide polymorphism analysis was conducted by whole-genome alignment against K12-MG1655 *E. coli* (wild-type laboratory strain) (39) and variant calling using Snippy Core v3.1 (https://github.com/tseemann/snippy). Phylogeny of core genomes was determined using IQ-TREE v.1.6.7 (substitution model: GTR + F + R10) with 1,000 bootstrap replicates. Phylogeny and metadata were visualized with Microreact (https://microreact.org/showcase). Branch lengths correspond to the number of SNPs difference (scale bar bottom left). MLST for each strain was determined using Enterobase (http://enterobase.warwick.ac.uk/species/index/ecoli). Asterisks indicate bootstrap support of 100% from 1,000 replicates.

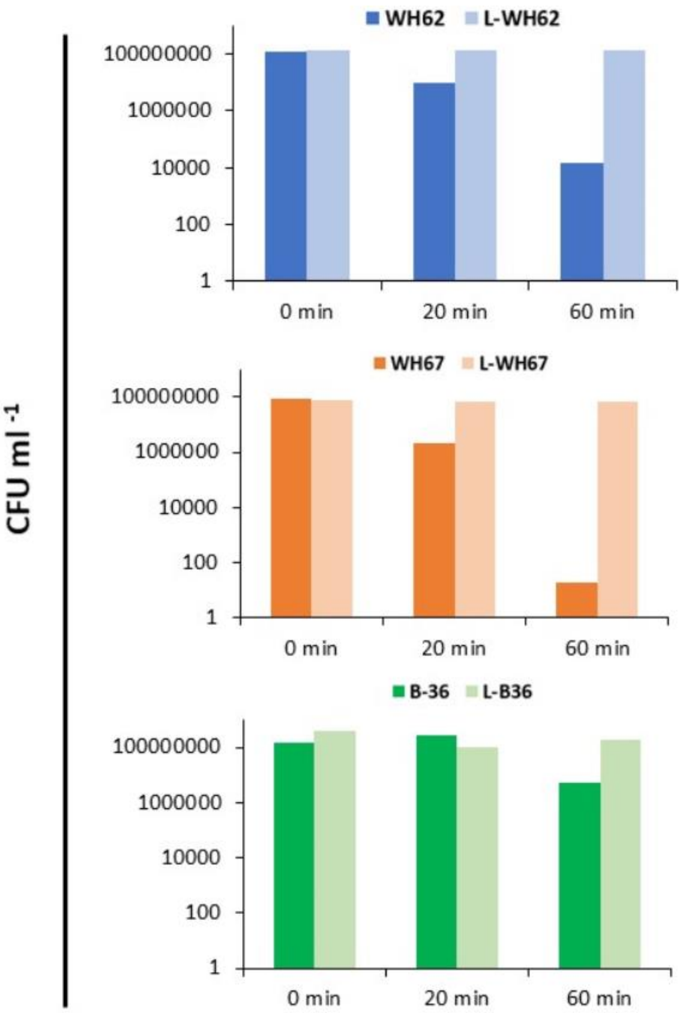


**Extended Figure 2.** Tolerance to sublethal oxidative stress (10 mM hydrogen-peroxide) in *E. coli* L-forms induced with meropenem.

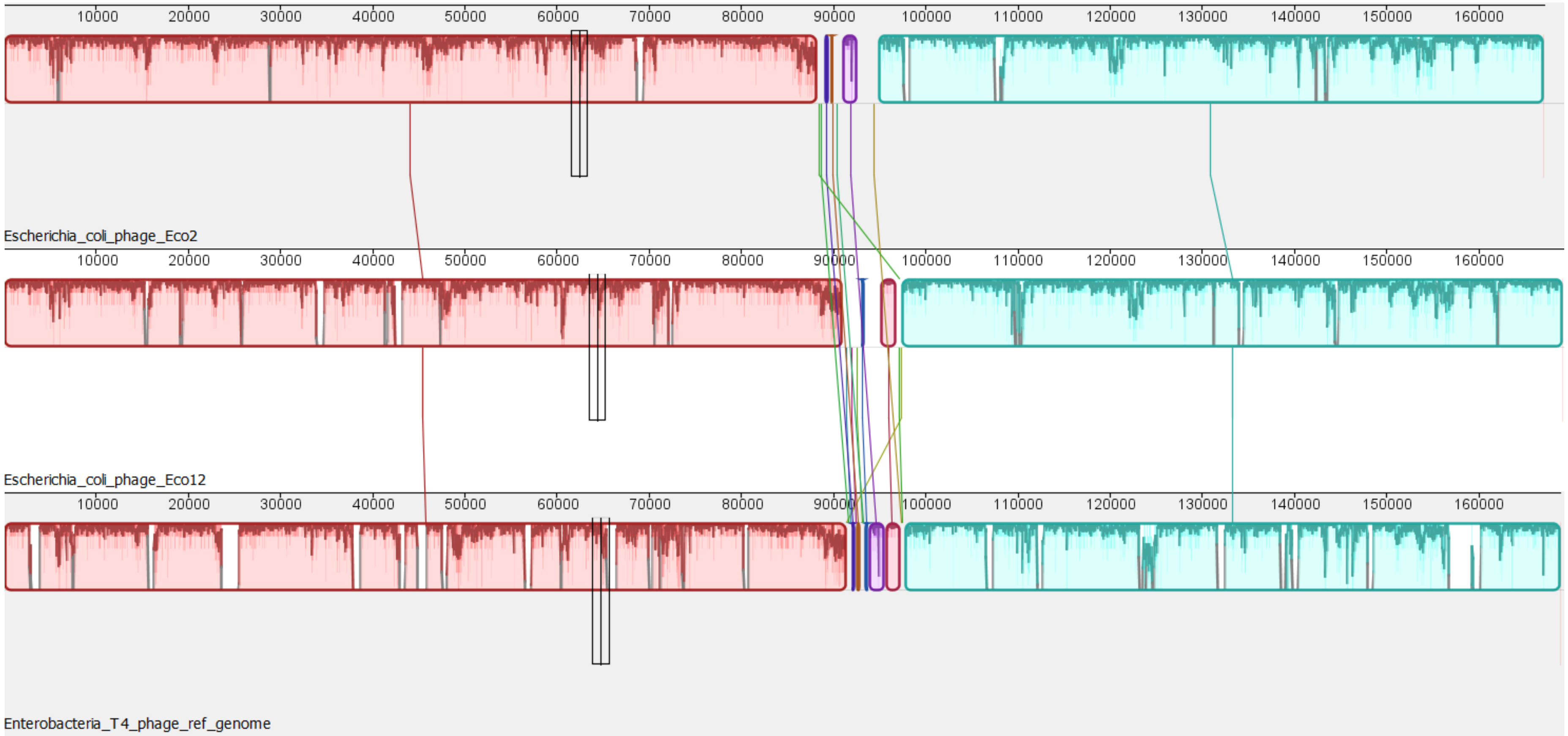


**Extended Figure 3**. Whole genome alignment with progressive MAUVE (34) of T4-like phages (*Myoviridae*). From top is Escherichia phage vB_EcoM_2, Escherichia vB_EcoM_12 and Escherichia T4 phage (reference; GeneBank NC_000866.4). The degree of DNA sequence similarity is indicated by the height of the red-coloured regions.

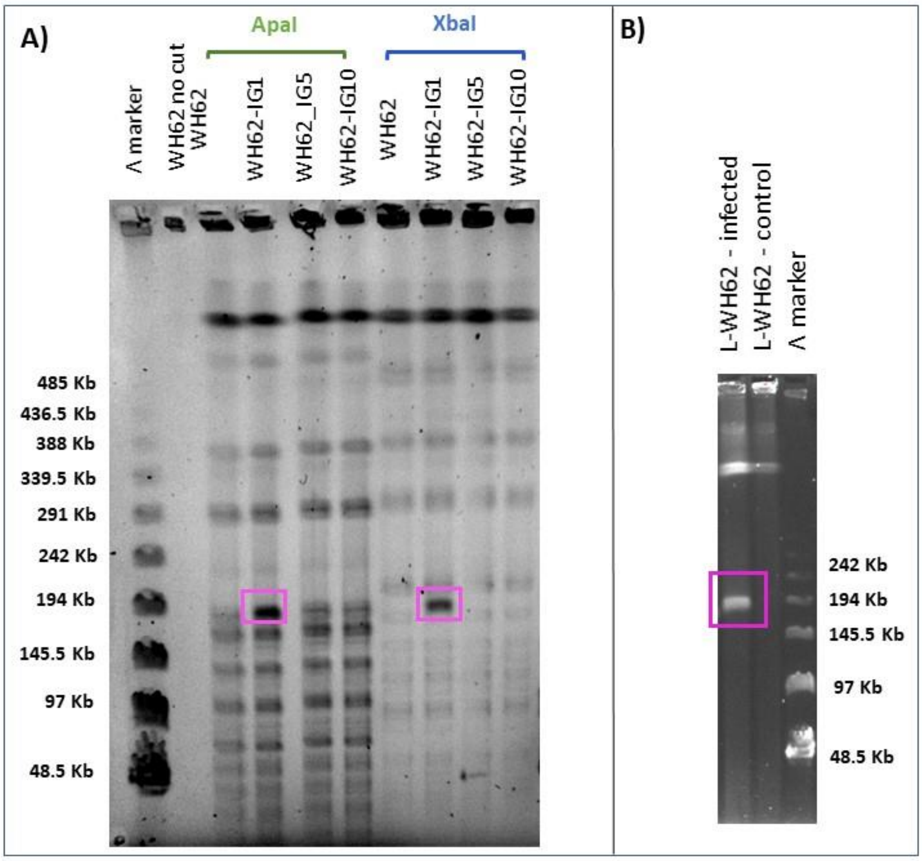


**Extended Figure 4.** PFGE of WH62 *E. coli* L-form infected (pseudolysogenised) with Eco12 phage. A) S1 enzymatic analysis of WH62 L-form pseudolysogen DNA digested with *Apa1* (sensitive to Dcm and *CpG* methylation) and *Xba1* (sensitive to Dam methylation) indicating presence of methylated episomal Eco12 phage (pink square; ~200Kb). B) PFGE of non-digested WH62 L-form pseudolysogen DNA indicating presence of ~200Kb episomal phage (pink square).

**Supplementary Table 1**. Primers used in this study for transcriptomic analysis of *E. coli* L-forms

| Primer | Sequence (5′-3′) | Target | Source |
|---|---|---|---|
| rpoB-F | GTAAGGCACAGTTCGGTGGT | *E. coli* rpoB for RT-PCR | Kamruzzaman et al, 2015 |
| rpoB-R | ATTTCCTGCAGGGTGTATGC | | |
| dinB-F | TTGCTTCCGGGGCGCTTTGA | *E. coli* dinB for RT-PCR | This study |
| dinB-R | ACGCCGTCAGTTGCAGCTCG | | |
| elaB-F | ACGACCTGACGCTGCTTAGT | *E. coli* elaB for RT-PCR | This study |
| elaB-R | AGCACGATAAACTGCCTGCT | | |
| oxyR-F | TGAGGTGAAAGTCCTTAAAGAGATG | *E. coli* oxyR for RT-PCR | This study |
| oxyR-R | GTCTGTGCTTCATGCAGATACATT | | |
| rpoS-F | GGCGTTGCTGGACCTTATCG | *E. coli* rpoS for RT-PCR | Kamruzzaman et al, 2019 |
| rpoS-R | TCAATCGTCTGGCGAATCCA | | |
| recA-F | TCCGGTAAAACCACGCTGAC | *E. coli* recA for RT-PCR | |
| recA-R | CGTGCGTAGATTGGGTCCAG | | |
| lexA-F | CGCGGCTGAAGAACATCTGA | *E. coli* lexA for RT-PCR | |
| lexA-R | GCGGCAACCCTTCTTCCTCT | | |
